# Evolution and favored change: a principle of least selection

**DOI:** 10.1101/2021.06.27.450095

**Authors:** Michael Yarus

**Affiliations:** Department of Molecular, Cellular and Developmental Biology, University of Colorado Boulder, Boulder, CO 80309-0347

**Keywords:** RNA world, replication, translation, genetic code, Normal distribution, truncation selection, fitness

## Abstract

Favored biological characteristics can evolve by a subtle path, but beneficial selection has predictable qualities to guide thought. These favored pathways are paths of “least selection”. Faster evolution is least selection, more probable because earlier evolutionary success is simply, “success”. A more likely path also requires least selection in the form of least selected change. Truncation selections, accepting only extreme values of a distributed quantity, produce greater change in population means. Truncation selection therefore readily offers a least selection. Assuming selection for a Normally-distributed quality, truncation is enhanced via simple dependences on increased standard deviation and higher selection threshold (consistent with some population survival). Least selection applies to both chemical and biological evolution, and can be estimated in general form, without reference to its genetics, from an underlying phenotypic distribution. Chemical truncation selection is free of the Haldane cost of natural selection; potentially yielding very rapid early evolution. Notably, a principle of least selection unifies prior examples of quick evolution. For example: the ‘crescendo’ of accurate codes leading to the Standard Genetic Code facilitates a least selection. More generally, evolutionary extinctions and radiations are events with multiple coercive thresholds; thus Earth’s history offers many wide-ranging truncation/least selection events.

## Introduction

### Evolution’s path can meander

Evolution readily takes a devious route. A natural population may include an opportune variant accidentally satisfying a proto-biological need. Those rare structures, perhaps less related to previous materials, can still provide the most likely evolutionary advance. Moreover, forms known from laboratory experiments are unlikely to completely sample possible protobiological conformations or chemistry: we are unsure of initial conditions. Finally, a conceptual barrier to study of evolution is that even the simplest replication, metabolism or expression are complex molecular events – thus, there will be no obvious primordial route to them. Research must therefore identify the *least improbable* routes to biological complexity.

Given these conceptual and experimental barriers, there is an acute need for well-founded predispositions about selected change. Is there a general way to anticipate favorable selection’s rate? Is there a coherent comparison that predicts probable evolutionary transitions? This text concerns these problems, always with ancient conditions, and particularly election of the Standard Genetic Code, in back-of-mind.

This text is meant to be semi-independently readable, starting with numerical examples (just below), then previous studies of replication, translation and genetic coding as illustrations of least selection (starting at **A principle of least selection**). Analytical background for all argument is in **Methods**.

## Results

### The Normal distribution as an example

To quantitate the fate of exceptional qualities *x, x* frequencies must be known. The Normal, or Gaussian, distribution (Methods) has a special status documented by the Central Limit Theorem (1). The theorem shows that any property which is the mean of random variables, which may individually be distributed in any way, nevertheless approaches Normal distribution behavior as the number of variables increase. Normal and near-Normal behavior is accordingly widespread, making it a cogent example.

### Selection acting on an example Normal distribution

Explicit quantitation begins with selection for *x*, denoted:

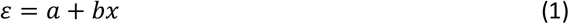

where *ε* is the number of descendants of individuals possessing *x*; *ε* is usually termed fitness. Within Eqn 1, *a* is increase independent of *x* quantity (for example, a concentration of *x*). Increase dependent on the advantageous quality is *bx*, with fitness proportional to *x*.

Fig. 1 shows unselected and selected distributions for *x*, under selection for fitness. The logarithmic vertical axis points up lesser change near the mean *μ* (including no change <u>at</u> *μ*), and resolves the selected increase in upper tail *x*. An initial Normal population (blue) is symmetrical around mean *μ* = 10. After selection for *x*, the selected, normalized population (Fig. 1, red dotted) has depressed *x* frequencies on the left, and elevated *x* frequencies above the mean on the right, required for selected frequencies to normalize to 1. *Δμ*_*fit*_ is the fractional increase in mean *x* due to fitness, *Δμ*_*trunc*_ the fractional increase in mean *x* due to truncation at *x*_*t*_. In Fig. 1, fitness selection increases *μ*_*fit*_ *by ≈* 4% (Me*t*hods Eqn 7; *Δμ*_*fit*_ = (σ*/μ*)^2^ ≅ 0.04; dashed red arrow, Fig. 1).

**Figure 1.**
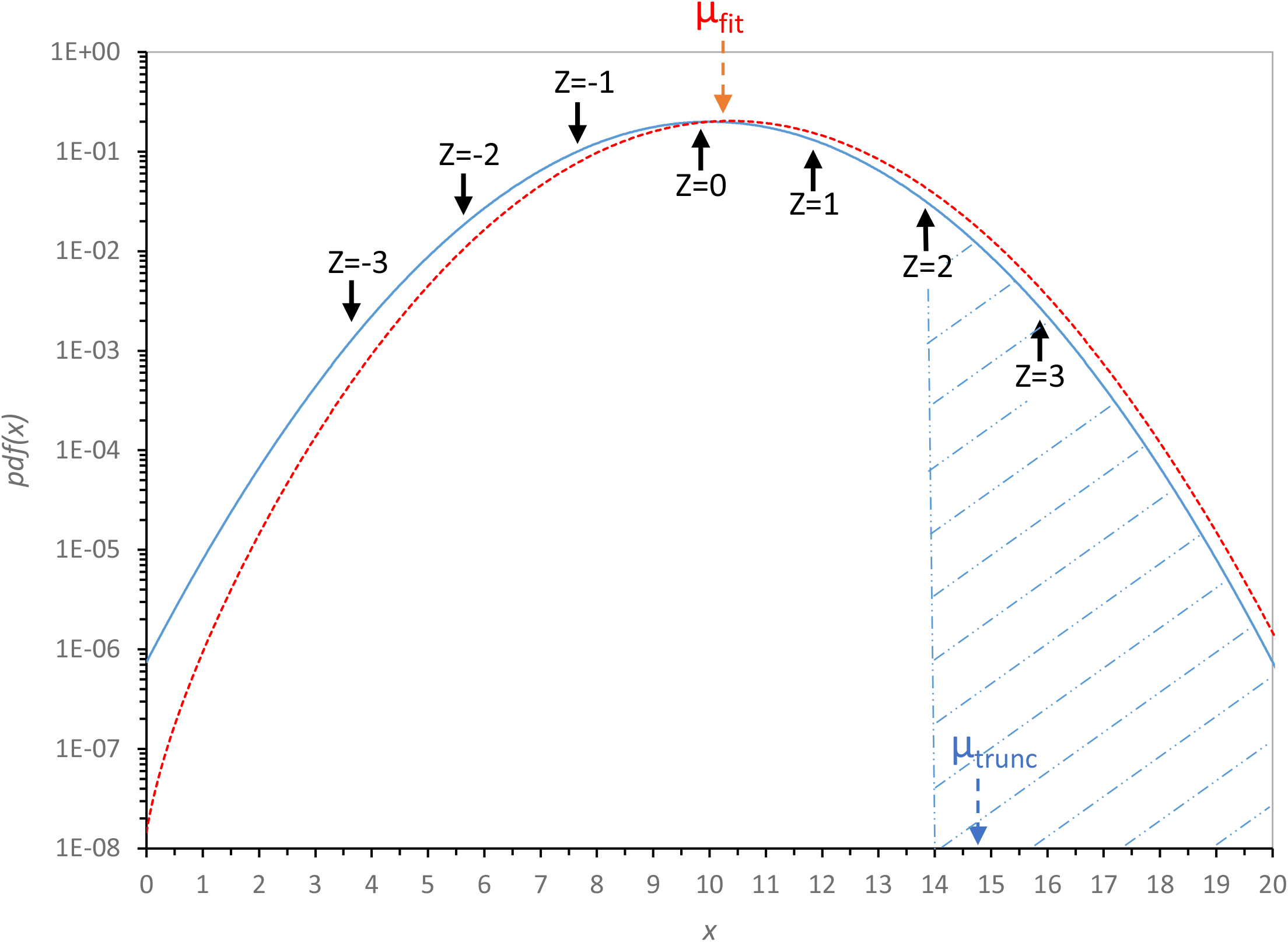
Fitness and threshold selection on a Normal distribution. The Normal probability of *x, pdf*(*x*), is plotted as a solid blue line against a logarithmic ordinate: in this example, mean *μ* = 10, standard deviation *σ* = 2. This distribution is selected, multiplying by *ε* = *a* + *bx* (dotted red line); *a* = 0.1, *b* = 0.5. A labeled red dashed arrow marks the resulting selected mean, *μ*_*fit*_ = 10.39, and black arrows illustrate the relation between the distributed quantity *x* and the generalized Z co-ordinate for the initial Normal. The dashed blue vertical line marks a selection threshold at *x*_*t*_ = 14, *Z*_*t*_ = 2, and its selected mean (*μ*_*trunc*_ = 14.746) is right-adjacent, and dashed blue. Visible threshold-selected area is hatched with light blue diagonal dash-and-dotted lines.

### Truncation selection

Evolution may also truncate a distribution, propagating a favored greater subset having more than some threshold *x*_*t*_ required to survive and reproduce (Fig. 1).

### *Z*_*t*_ generalizes truncation discussion

Description of truncation is facilitated by Z co-ordinates. *Z* values are distances from a mean in σ units:

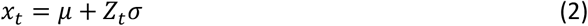

*Z* and *Z*_*t*_ (the truncation threshold) can also be negative, measuring distance below the mean (Fig. 1). Use of *Z* eases comparisons because standardized Normal distributions all have the same probability at the same *Z* (Methods). Φ(*x*_*t*_) and Φ(*Z*_*t*_) reference the same *x* and are unaltered by switching to *Z* (as are *ϕ* and *ϕ/*Φ), but *x*_*t*_ and *Z*_*t*_ are distinct (Eqn 2).

### Truncation and fitness selection compared

Fig. 1 previews a general conclusion; truncation is usually more effective; in fact, *Δμ*_*trunc*_*/Δμ*_*fit*_ *>* 12 for this Fig. 1 example, comparing downward red and blue dashed arrows for selected *μ*.

In greater detail: Fig. 2A shows that here, a small *ε* effect quickly reaches a maximum *Δμ*_*fit*_ = 0.04 (blue data, *σ* = 2) as *b* (and the selective effect of *x*) increases. Non-linear improvement of *Δμ*_*fit*_ with *σ*^2^ is also shown by results for varied *σ* values (Methods Eqn 7; dotted and solid lines are explained below).

**Figure 2A.**
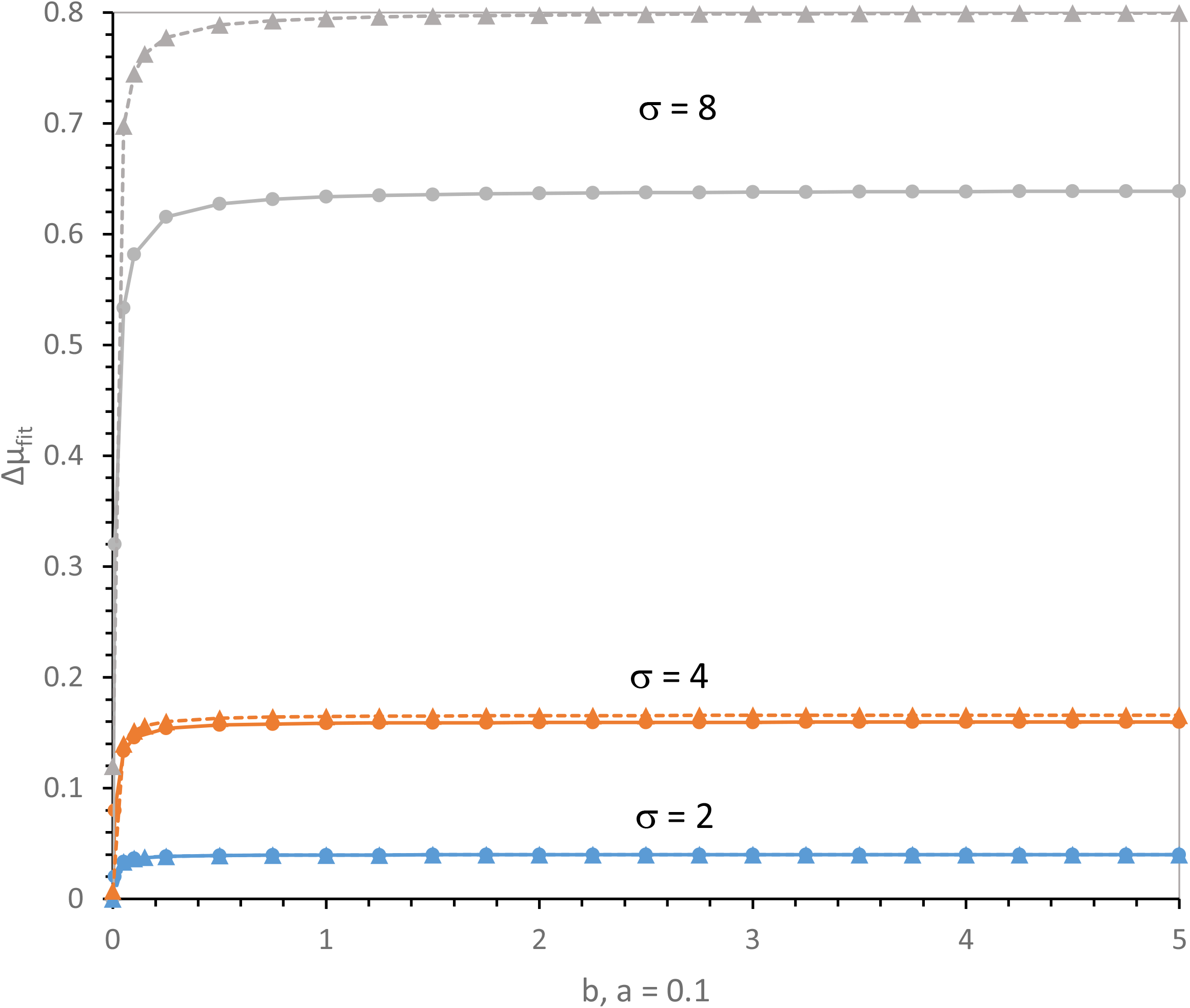
A. Selected fitness increase in mean *x, Δμ*_*fit*_, with varied dependence on *x*. *ε* = *a* + *bx*, with a = 0.1 and b varying. Normal distribution parameters are those in Fig. 1. Solid lines are calculated for full distributions (Methods Eqn 8), dotted lines for truncation of broad fitness distributions at *x* = 0 (Methods Eqn 14). The lines for *σ* = 2 are too similar to be resolved.

**Figure 2B.**
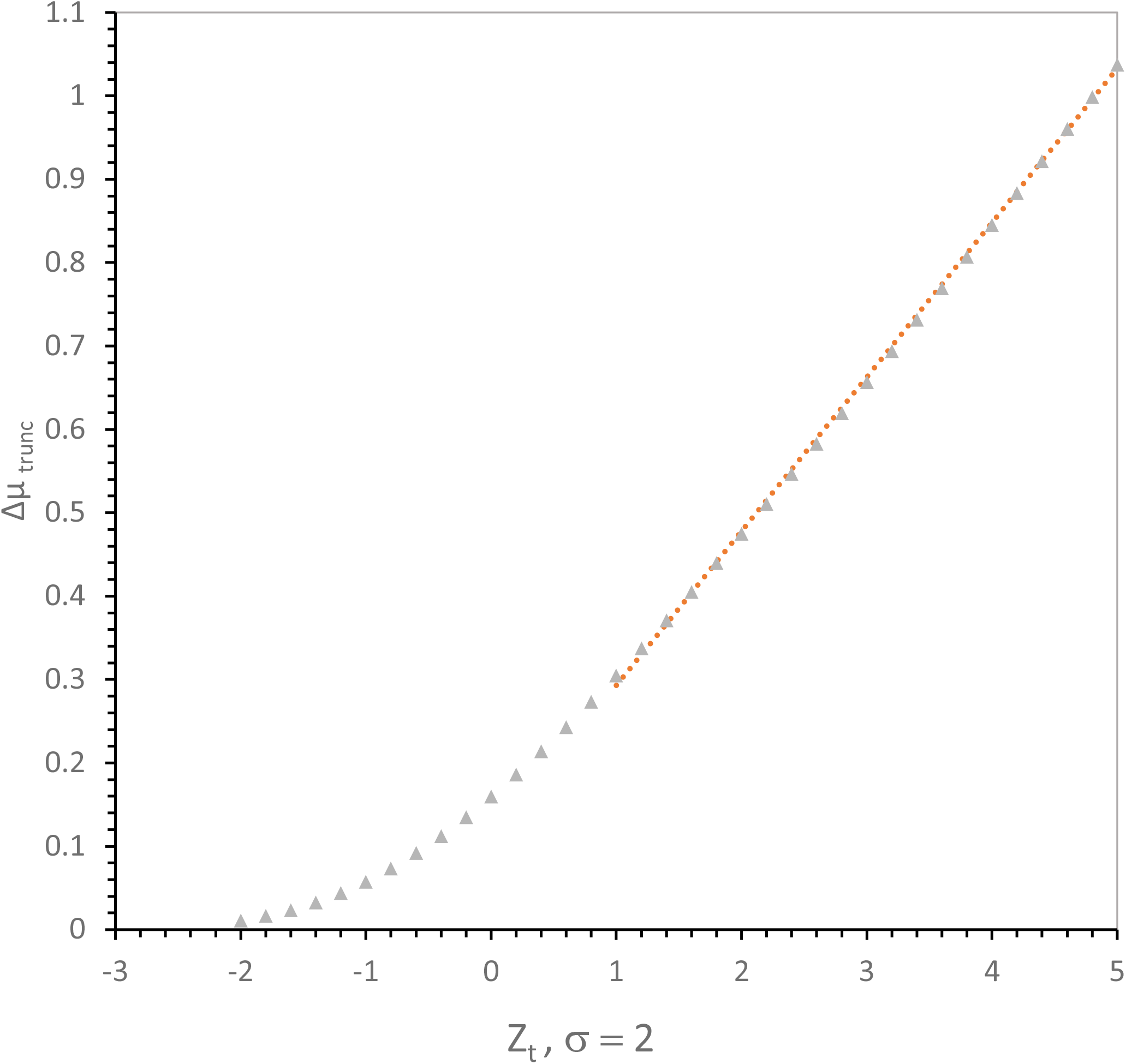
Selected truncation increase, *Δμ*_*trunc*_, across a Normal distribution as the acceptable threshold *Z*_*t*_ varies. *Z*_*t*_ ranges from below the mean (*Z*_*t*_ = −3) to far above (*Z*_*t*_ = 5). An arbitrary linear red dotted line has been superposed on data to emphasize linearity at *Z*_*t*_ *≥* 1. Normal parameters are those in Fig. 1.

In Fig. 2B, truncation behaves differently: as *Z*_*t*_ approaches the mean, *Δμ*_*trunc*_ becomes increasingly significant, and ultimately increases steeply and linearly. Notably, with example values, a threshold *Z*_*t*_ *≥* 4.8 (Fig. 2B) more than doubles initial *μ* (*Δμ*_*trunc*_ *≥* 1). Substantial evolutionary change occurs via one truncation, in contrast to fitness selection (Fig. 2A). The superiority of truncation selection is retained even if *x* truncation is gradual rather than sharp (2,3). Accordingly, threshold selection has a special role in biological change.

As the apposed straight (dotted, red) line in Fig. 2B indicates, truncation selections (*Δμ*_*trunc*_ and *Δμ*_*fit, trunc*_ below) improve *≈* linearly above *Z*_*t*_ = 1 (above *≈* 0.3-fold selected increase).

### Effect of σ on evolutionary change

Fig. 3A shows how initial variability (as standard deviation) *σ* alters selected improvement, *Δμ*. Different selections exploit initial variation *σ* with different efficiencies.

**Figure 3A.**
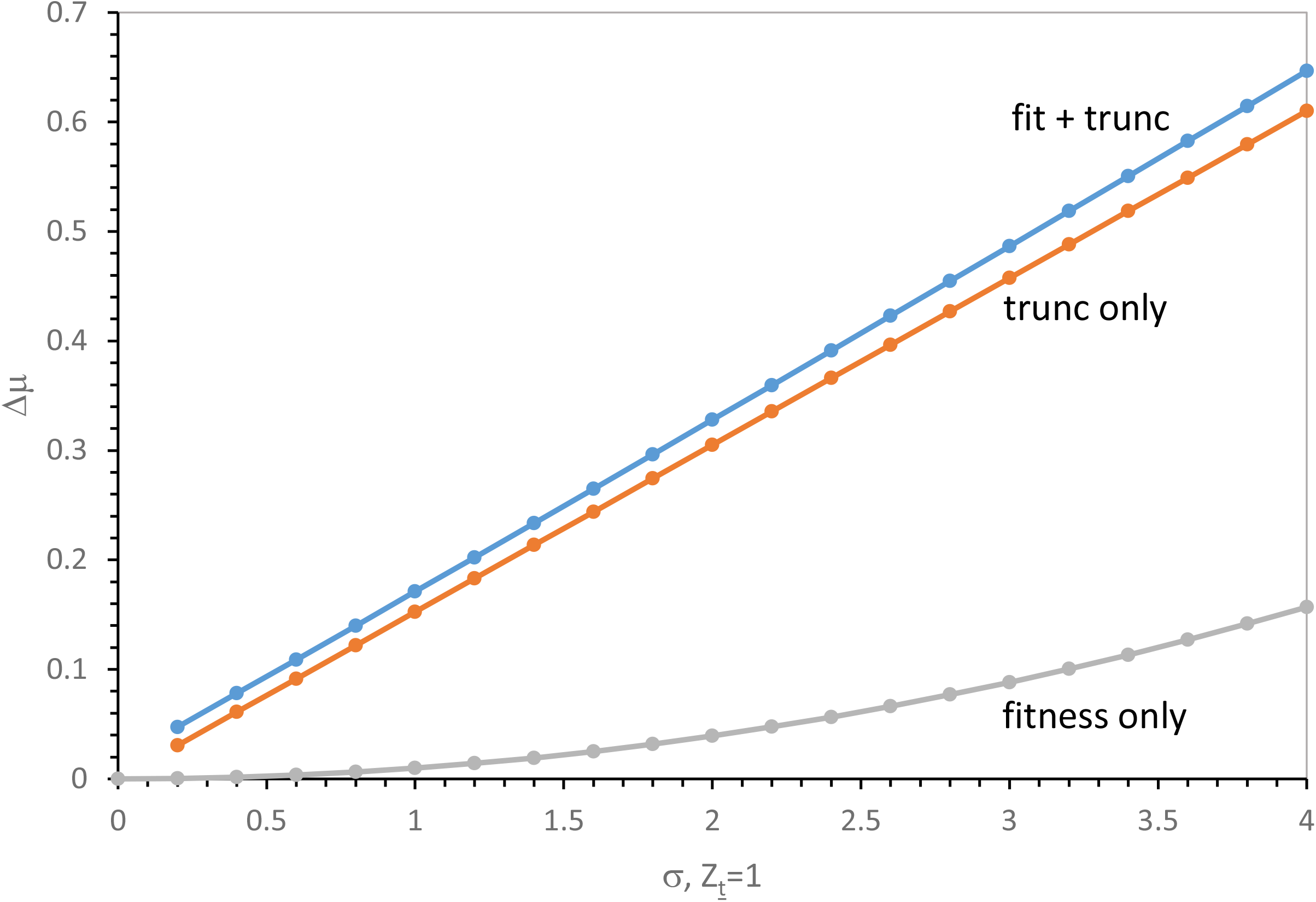
Selected increase in mean x,, *Δμ*_*fit*_, *Δ*μ_*trunc*_, *Δμ*_*fit, trunc*_ as a function of increasing initial standard deviation σ. This example assumes *Z*_*t*_ = 1 throughout. Other Normal parameters are those in Fig. 1.

**Figure 3B.**
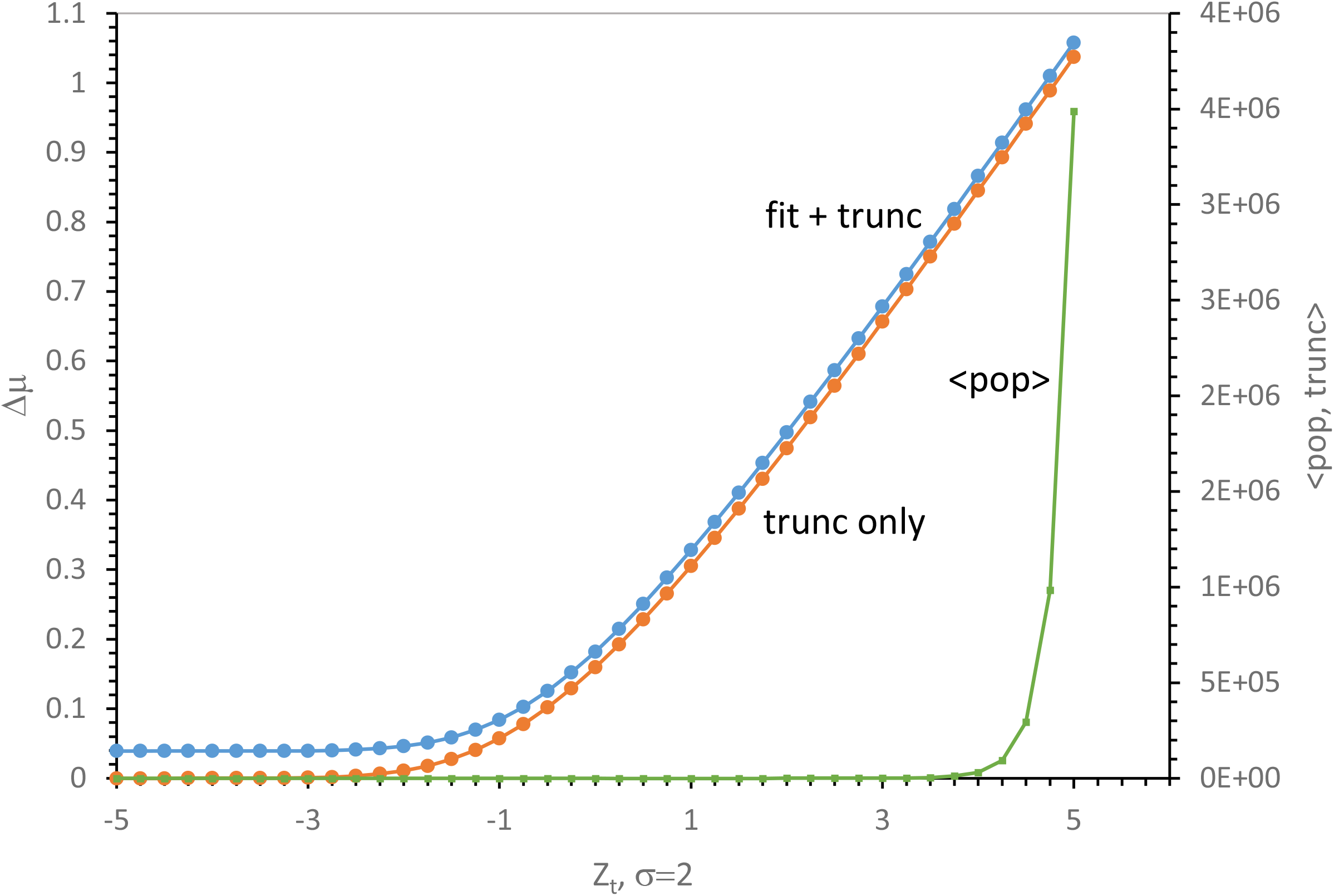
Selected increase in threshold-selected means, *Δ*μ_*trunc*_ and *Δμ*_*fit, trunc*_ as a function of threshold Z_t_. Standard deviation is **σ** = 2. The righthand ordinate is the population size required to find one improved individual above threshold *Z*_*t*_, using truncation alone. Other Normal parameters are those in Fig. 1.

Selected *Δμ* _*trunc*_ increases precisely linearly with *σ* (at constant *Z*_*t*_) with no selection possible if there is no variability (Methods Eqn 11; Fig. 3A).

Calculation of *Δμ*_*fit, trunc*_ shows that combining fitness with truncation (Fig. 1) adds a few percent to the superior effect of truncation on selection. *Δμ*_*fit*_ is generally less effective than either truncation, even given its benefit from larger initial *σ* (Methods Eqn 6, Fig. 2A). Most effective selection is via truncation or truncation+fitness, slightly be*t*er for both (Fig. 3B).

### Fitness with increased *σ* creates truncations

*Δμ*_*fit*_ responds to *σ*^2^ (Methods Eqn 7) and *Δμ*_*trunc*_ to *σ* (Methods Eqn 11), thus decreasing the gap between fitness and truncation selections as *σ* increases (Fig. 3A). However, increased variability *σ* does not yield indefinitely effective fitness selection. Large *σ* broadens distributions, so that individuals with *x* = 0 will exist. For usual qualities of evolutionary interest (concentration, height, weight, color…), *x <* 0 is often not meaningful. Thus, enhancing *Δμ*_*fit*_ by increasing *σ* creates truncations at *x* = 0 with be*t*er *Δμ*_*fit, trunc*_. This is shown in Fig. 2A, where dotted curves include the effect of truncation at *x* = 0 due to broad distributions (Methods Eqn 14). For this reason, and given the tendency to small fitness selection (Fig. 2A, 3A), over a large range of conditions, truncations yield faster evolution.

### Effect of threshold *Z*_*t*_ on evolutionary change

The above conclusion is reinforced by the effect of varied thresholds (Fig. 3B). Here fractional selected mean increase, *Δμ*, is presented versus varied threshold *Z*_*t*_ (constant standard deviation, *σ* = 2). *Δμ*_*trunc*_ is small at *Z*_*t*_ *<* −2, but then rapidly becomes significant, decisively increasing above the mean.

With simultaneous fitness and truncation, *Δμ*_*fit, trunc*_ is significantly increased even at *Z*_*t*_ = −5 because it includes an intrinsic small but complete fitness selection (Methods Eqn 8). However, such fitness is superior to truncation only far below *μ*; truncation becomes the major effect at *Z* just below the mean (Fig. 3B).

Because of rapid decline in Normal *pdf(x)* at higher *Z*_*t*_ (Fig. 1), linear increase in *Δ*μ_*trunc*_ and *Δμ*_*fit, trunc*_ at high *Z*_*t*_ (Fig. 3B) are explicable. *Δ*μ_*trunc*_ approaches *Z*_*t*_ as *Z*_*t*_ increases because selected individuals above *Z*_*t*_ become increasingly close to *Z*_*t*_, assuring similar behavior for *Z*_*t*_ and *Δμ*. Further, comparing the strict limit for *Δμ*_*fit*_ in Fig. 2B with this unlimited linear increase in *Δ*μ_*trunc*_ and *Δμ*_*fit, trunc*_ (Fig. 3B), truncation superiority is reinforced.

### The cost of truncation

Truncation selection at greater *Z*_*t*_ is strongly constrained by the rarity of exceptional *x*: population size required to expect one improved individual above *Z*_*t*_ is shown on the righthand axis in Fig. 3B. Normal tails decline increasingly steeply with *Z* (Fig. 1), and the population needed to recover one improved *x* individual increases steeply and non-linearly with *Δμ*. In Fig. 4, the population expected to contain one selected individual, 1*/*Φ(*Z*_*t*_), is compared to selected improvement *Δμ*_*tfit*_ for different initial variabilities, σ.

**Figure 4.**
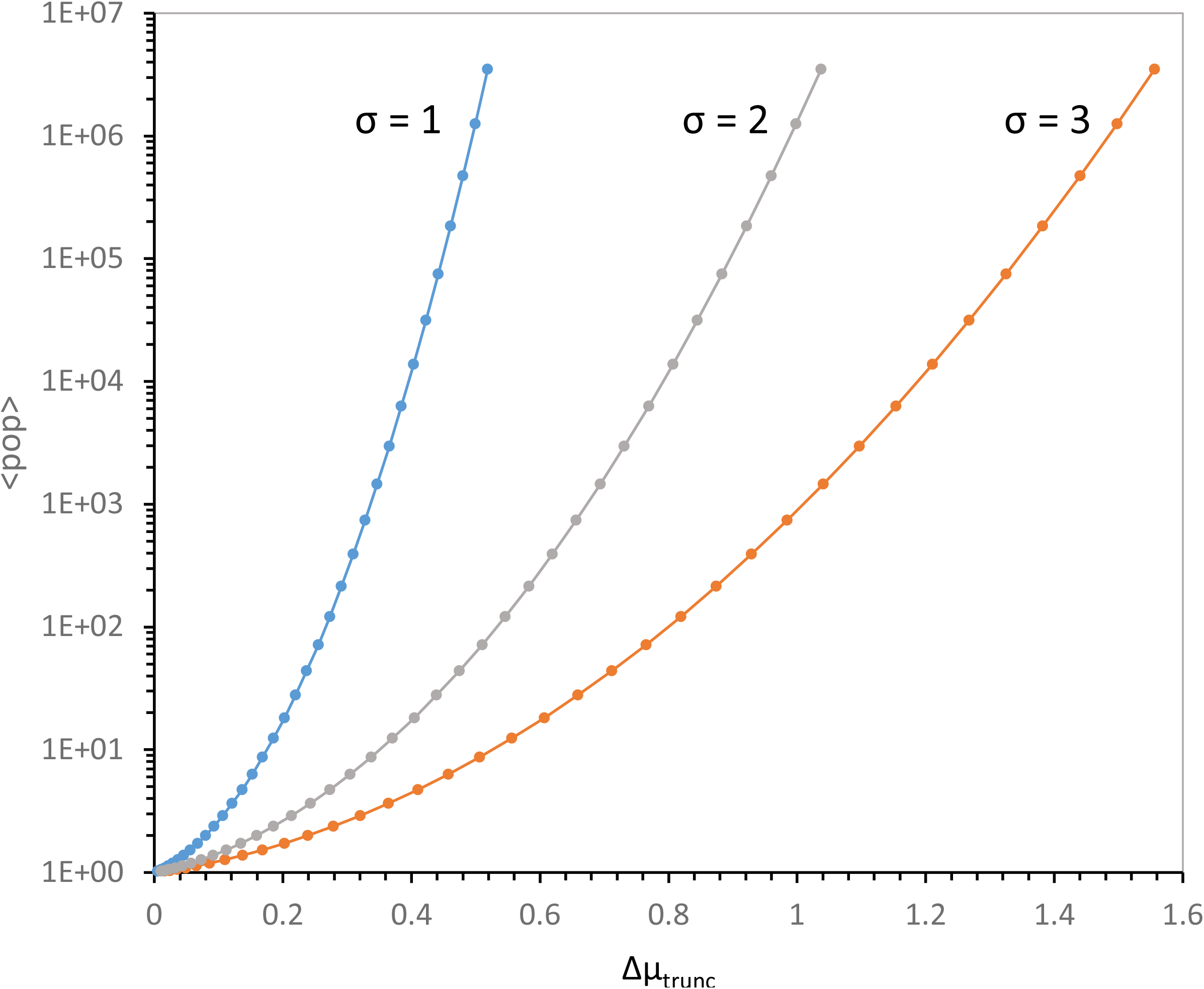
Population sizes to recover one superior individual as a function of selected improvement, Δμ_*trunc*_. Population sizes, 1*/*Φ(*Z*_*t*_), are plotted on a logarithmic ordinate, assuming different starting variabilities, *σ* (Methods Eqn 11). Normal parameters are those in Fig. 1.

Substantial improvements (e.g., 50% of *μ, Δμ*_*trunc*_ = 0.5) can be selected in populations with varied standard deviations. But such selection is critically dependent on initial variation - decreasing initial *σ* 3-fold increases the population required more than 5 orders of magnitude (Fig. 4). So, selection of an upper tail for moderate improvements is undemanding. But, if exceptional increases are vital – much larger populations, low survivals, and/or elevated variabilities (*σ* or *σ/μ*) are required. This constraint can be severe, with near-exponential population increases required for small improvements. Accordingly, finely divided populations with small individuals, unusually varied populations, or both were surely crucial to rapid chemical evolution. Varied microbes probably executed Earth’s emergent biology (Fig. 4).

### Chemical selection is not slowed by the cost of natural selection

The cost discussion above has a passing resemblance to the classical discussion of the cost of natural selection, due to JBS Haldane (4). However, biological and chemical cases differ greatly. Haldane’s cost was due to replacement of genes in a modern, genetic population: possessors of an old gene configuration must perish to be replaced by a new genetic constellation. Too much replacement yields excessive perishing, and thus extinction. Haldane’s cost may be mitigated by recombination (5), so may help account for the prevalence of sexual reproduction. Still, the rate of <u>natural</u> selection is cost-limited. However, in a non-genetic population, any fraction of a population may be replaced in a short time if a small selected portion persists and reproduces. Thus, chemical least selection may be sporadic, but much more rapid than natural selection. As previously calculated, inheritance itself can emerge in a few days (6).

### A principle of least selection

A particularly useful idea arises from minimizing time-to-select. Maximal improvement *Δμ* minimizes selections and thus minimizes time-to-evolve a particular *x*, thereby also indicating the most likely evolutionary route. Minimal time and least selected change required can accommodate multiple evolutionary effects, incorporating both threshold and truncation, evoking the faintly parallel physical principle of least action (7). Finding this extremum identifies the most likely route to an improbable outcome. Call this joint minimization of selections or time-to-select “least selection”. Next, the explanatory value of a principle of least selection is illustrated, using prior molecular biology examples.

## Discussion

### Outline

Fluctuation frequently is the crux of evolutionary progress (8). This makes some evolutionary progress predictable (9). Here, this idea is quantified: functions of standard deviation *σ*, coefficients of dispersion *σ/μ*, threshold *Z*_*t*_, and/or selection parameters *a* and *b*, numerically govern selection’s Normal effectiveness. The results, viewed broadly, are simple.

For Normally-distributed *x*, truncating by requiring a threshold *x*_*t*_ yields maximal increase in a favored quality. Truncation-selected change is usually greater than from selection of fitness, and progress is proportional to standard deviation *σ* or dispersion *σ/μ*. Truncation’s Improvement in a favored property also is proportionate to an acceptable *x* threshold, *Z*_*t*_, at larger *Z*_*t*_. However, threshold selection is effective even at *Z*_*t*_ = 0; such a threshold at the mean sums a region of positive effects. Populations with highest *σ* and truncation threshold *Z*_*t*_ require least selection, and evolve fastest. However, exploiting upper-tail *x* for large change requires large populations (Fig. 2B, 4).

### Here, selection applies only to phenotypes

Present calculations follow distribution properties, thus conclusions do not reflect particular system features. Instead, only the initial and final means of expressed qualities, on which selection acts, matter. An assumed genetic system is required to deduce selection’s genetic consequences. But ignoring genetics has a strategic advantage: present selection quantitation applies to any genetic system whatever.

A phenotypic focus therefore cuts two ways; we cannot find out anything about genetic systems, without further assumptions, and we assume that changes are substantially heritable.

However, results apply to any Normally-distributed property, seemingly particularly appropriate to quickly varying primordial, pre-Darwinian systems.

### Example: Starting bloc selection exploits an inevitable threshold

The starting bloc (10) was encountered while calculating that known ribonucleotide reaction rates suffice for evolution of encoded, and therefore heritable, chemical capacities (6).

Starting bloc heritability depended on two well-known RNA capabilities. The first is existence of simple catalytic RNAs, homologues of the adenylate-dinucleotide-containing enzyme cofactors (11,12). The second is observed templated synthesis of similar molecules via an ultimately simple, geochemically plausible (13) RNA “gene”, poly-U (14). Selection of such poly U-templated cofactor congeners (in preference to untemplated chemical synthesis) is uniquely effective early, when only some sites have begun synthesis – these early-acting individuals are the “starting bloc” (10). Increases of templated synthesis selected in such a population are nearly proportionate to dispersion (*σ/μ*) in the chemically active product over a large range of product concentrations (10), just as calculated here for least selection (Fig. 3A).

### Starting bloc selection employs distribution tails

The original starting bloc produced roughly proportionately faster change when carried out with lower survival after selection, a condition there called “stringency”. This parallels present calculations: increased threshold *Z*_*t*_ and decreased survival in Fig. 2B, 3B produce faster selection. Similarly, combining variability (as in dispersion *σ/μ* or standard deviation *σ*) with survival in a single index best predicted starting bloc success (10).

### Starting bloc selection exploits an early threshold

The difference between starting bloc loci that make an active molecule early and inactive ones with no product is particularly readily selected. Starting bloc success via some, versus no, early activity exploits truncation, paralleling least selection in Fig. 3B. Accordingly, selection of encoded chemical capacity in a starting bloc (6,10) follows, with almost formulaic precision, the variability, survival and threshold dependence for least selection (Fig. 3A, 3B, 4).

Starting blocs may be of fundamental evolutionary significance: nascent biological capabilities are intrinsically selectable. In particular, encoded (genetic) expression is a crucial biotic innovation. While replication is often cited as the foundational requirement for biology (15), there seems small advantage in replicating unexpressed sequence information. Thus, effective expression likely predates replication (12). Accordingly, a starting bloc’s least selection potentially speeds expression united by a genome, thereby initiating vertical inheritance and Darwinian evolution.

### Example: Chance utility captures favorable variation

Prebiotic systems must evolve toward biological complexity before genomes, or genome-directed biology would be inaccessible. Selected temporary, or especially, permanent change in chemical systems before the advent of genetic inheritance has been called ‘chance utility’. For example, variation in a 100-fold more concentrated inhibitor frequently provides an opportunity to capture a useful, but apparently overwhelmed minor reaction (16). This accidental contravariation of a substrate and its competitor relies on favorable *μ* and *σ* for these reactants; that is, on chance resemblance to a favorable result, an opportunity for least selection.

### Example: Random reactant supplies necessarily support near-ideal reactions

Even given random, uncontrolled nucleotide supplies, a hypothetical small RNA replication recurs (8). Notably, most RNA replication in the randomly-fed reactor comes from a recurring subset of near-ideal substrate arrivals. So, a randomly-fed pool produces recurrent specific results from u*t*erly unregulated inputs, an opportunity for least selection.

### Least selection opportunities can flicker

The starting bloc, chance utility example, and recurring near-ideal reactions illustrate a common theme: selection can exploit kinetics. Opportunities for least selection can flicker: transiently appearing, though usually absent. Truncation exploits favorable transients; thus relatively rapid selection in a world that varies more slowly greatly enhances truncation and consequent least selection.

### Example: Stable optima identify evolutionary paths

Just as flickering opportunities for truncation predict evolutionary transitions, stable optima identify persistent probable paths. An example is advent of initial, simplified wobble coding. In the forming genetic code, there appears an optimal late time for wobble advent (17). A time of maximal SGC resemblance offers a least selection, marking that era as plausible for wobble’s rise.

### Example: Bayesian convergence follows from least selection’s extremum

Bayes’ theorem says that the best origin hypothesis explains all aspects of a phenomenon. However, an explanation is uniquely effective if independent events are explained, and the hypothesis’ probability is elevated by the product of independent confirmations (18). For example, a fusion route to the Standard Genetic Code (19) explains how metabolic, chemical and wobble order coexist without mutual interference in a near-universal coding table. Bayesian convergence is robustly supported by least selection, which maintains that the historical route to the genetic code was distinct, perhaps unique, because it traverses a continuous extremum. This in turn overlaps Orgel’s dictum that evolution exhibits Continuity (20); that is, later evolutionary states are necessarily linked to their precursors by continuous change.

### Example: Distribution fitness is least selection, using different language

The fitness of a superior group within a distribution is a key to apparently improbable outcomes. For example, a small minority of nascent coding tables has unlikely coding properties (Table 2; Ref. 9). These become ordered quickly, having made accurate Standard Genetic Code (SGC) assignments (9), but leaving some unassigned triplets that will later be critical during final assembly of a complete code (19). Optimal tables appear quickly by avoiding slow events, like assignment decays (9). This useful minority of genetic codes short-circuits the outward problem of evolving a complex pattern by accidently evolving it simply. Apparently implausible behavior; for example, showing what seems to be foreknowledge of late coding early in code history, is possible because they are a rare minority selected late. Their properties ultimately seem near-ideal, but pre-exist without purpose. When selected as most functional among a population (21), they immortalize their improbable histories, thus presenting a seeming paradox. This peculiar quality is routine in truncation selections, which by definition capture improbable states (Fig. 1, 4).

Thus, distribution fitness anticipates least selection - success from rare precursors with unexpectedly close resemblance(s) to an upcoming biological achievement.

More recent work on coding (21) clarifies the search for excellent minorities. Fusion of early partial codes strongly enforces code uniformity, because fused unlike codes assign ambiguous codon meanings, and are less fit. Fusion therefore generates a long-lived series of codes that increasingly resemble an underlying consensus, such as the Standard Genetic Code. Given a progressive increase in code uniformity (called the ‘crescendo’) that presents a long series of varied SGC-like encodings, code fusion proficiently implements least selection of the SGC (22).

### Example: Least selection appears as similarity to archaic forms

Vast gaps in time can link similar things, implementing the “least” in least selection. Ribosomal peptidyl transferase is an example. CCdApPuro (23) is a rationally-designed oligonucleotide inhibitor that binds both halves of the ribosomal active site, contacting A- and P-sites in a configuration that allows peptide transfer between them. So, CCdApPuro emulates the transition state for peptide synthesis (24). An RNA selected from randomized sequences to bind CCdApPuro with affinity similar to ribosomes (25) exhibited two families of binding sites with 17 nucleotide matches to the peptidyl transferase loop of rRNA. This homology included an exact 8-mer match with among the most conserved of all RNA sequences, the ribosomal peptide transferase active site AUAACAGG. CCdApPuro also bound this RNA octamer in the large ribosomal subunit (26), in an orientation similar to selected RNA sequences (25,27). Selected sequences even included a likely catalytic feature, an A with a shifted pKa, though catalysis was not selected (27). These results support initial peptide synthesis in an RNA cradle, then least selection of a ribonucleoprotein descendant, still resembling its RNA ancestor billions of years later.

### Truncation is a major evolutionary event: extinctions

We know macroscopic evolution mostly from durable fossils of relatively modern, multicellular creatures. Such evolution is sporadic, with long periods of relatively slower change interrupted by shorter eras containing mass extinctions (28,29). There have been *≥* 7 relatively recent, major extinctions. Hodgskiss et al (30) present isotopic evidence for one such ending the Great Oxidation Event 2 gigayears ago. For the last 540 million years, see Muscente et al (31) for such alternation of stability and catastrophe in the “big five” extinctions (32). Ceballos (33) provide evidence of a current, human-induced catastrophe. More evident quick species losses are followed by re-diversification of surviving organisms (34). Thus major extinctions are also radical multiple truncations, making least selection unmistakably relevant to life’s history on Earth.

### Truncation is a major evolutionary event: radiations

The mostly sessile oceanic life of the Edicaran era (588 - 539 Mya) had made major progress: it included metazoans who were bilateral, for example (35). However, these benthic creatures were almost completely replaced in a ‘Cambrian explosion’ (539 – 488 Mya) of novelty (36), in which most modern phyla were established (37,38). Exceptionally-preserved worldwide Cambrian fossils in Canadian Burgess shale (39) and at Qingjiang in Southern China (40) validate Cambrian radiation at the consensus origin of modern biota. Radiations, however radical, do not re-originate life. Thus major radiations are also radical multiple truncations, making least selection unmistakably significant to life’s history on Earth.

### Plausible biological change: least selections and biological intermediates

Seeking least selection will not suddenly manifest biotic history. For example, neutral and near-neutral change (41) have not been considered.

Even more significantly, plausible precursors may not yet be known, because our present molecular portfolio is deeply partial. For example, prebiotic nucleotide activation is an indispensable, but elusive missing link to both ancient translation and replication. Imidazole activates AMP for templated oligoRNA synthesis (42). Existence of a possible prebiotic, UV-activated imidazole activation of AMP (43) has guided related chemistry with 2-amino-imidazole and ensuing templated RNA synthesis from activated CMP and GMP (44). Thus, a vital activation gap may ultimately be filled - least selection will compare alternative solutions.

Least selection proposes a frequently-practical estimate of evolution’s path. It overlaps the suggestion that the most plausible route yields a known intermediate with greatest probability (9). Assembling least selections to plausibly intersect a known biological result creates a credible evolutionary conduit. When that evolutionary change can be reduced to quantitative alterations in selected characters, one will likely see regions with slower change (fitness selections, Fig. 2A, 3A) interspersed with change an order-or-more steeper (truncations, Fig. 3A).

## Acknowledgement

A preprint of this discussion is available (22).

## Methods

### Selection from a varied population

The probability of *x* within its population is often describable by a normalized ***p***robability ***d***istribution ***f***unction, *pdf*:

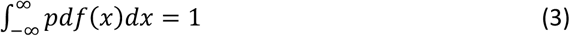

To follow a population’s fate, average *x, μ*, is followed. If *pdf*(*x*) is the probability of *x*:

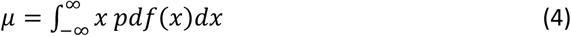

Selection changes *pdf*(*x*):

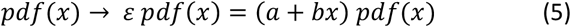

### Selection for fitness

Consider selected increase of a favored quality *x*; such selected increase in mean *x* is described by the Price equation (45). The Price equation mathematically partitions evolutionary change into that due to selection and that due to all other frequency changes for qualities of interest (46). We are particularly interested in selection:

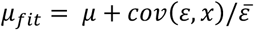

Where *cov* is covariance and 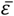 is population mean fitness:

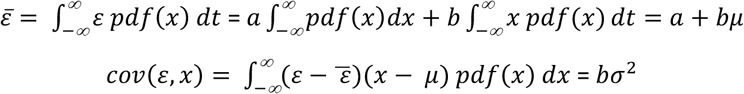

Thus

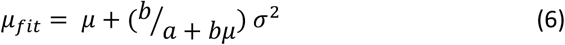

Fractional selected change in mean *x* is:

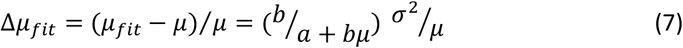

*Δμ*_*fit*_ defines selected improvement. *Δμ* is normalized to the initial mean, reducing dependence on the choice of scale. *Δμ* = 0 for no progress, and, scaled to *μ, Δμ* increases as evolutionary change increases. For example, *Δμ*_*fit*_ = 1 is selected mean doubling.

### Fitness entirely dependent on *x* is particularly simple

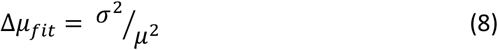

This plausible situation is emphasized in Fig. 2: selection requiring *x* and ∝ *x* (*a* = 0) increases average *x* precisely by coefficient of dispersion squared, (*σ/μ*)^2^, independent of *b. Δμ*_*fit*_ is wholly defined by the initial distribution’s shape. As intuition suggests, complete dependence on *x* (*a* = 0) yields maximal fitness selection; larger by the factor (*bμ* + *a*)*/bμ*, thus necessarily the best selection. The Price equation is a mathematical identity (46) and so always valid. Further, no specific system property is assumed in calculations. Thus Methods Eqn 6, 7 are true of any integrable *pdf*(*x*), thus any well-behaved distribution. *Δμ*_*fit*_ consequently describes genetic, non-genetic, prebiotic or solely chemical systems. Blue (before) and dotted red lines (after selection) in Fig. 1 illustrate fitness selection, using a Normal distribution example.

### Calculating the fate of Normally-distributed qualities

Because of the broad application of the Normal distribution, its integrals are evaluated in standard software. For example,

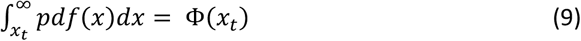

where Φ*(x*_*t*_) from spreadsheet or mathematical software is the population fraction exceeding a threshold: e.g., for *x* between x_t_ and ∞ (Methods).

Mean *x* for selection above a threshold, *μ*_*trunc*_, measures selection’s success, and can be evaluated by rearrangement:

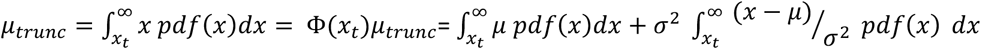

where Φ*(x*_*t*_) is defined by Eqn 8, the integral-normalizing factor for *μ*_*trunc*_. The rearrangement on the right can be integrated, because the second rightward term integrates the negative derivative of the Normal *pdf*(*x*):

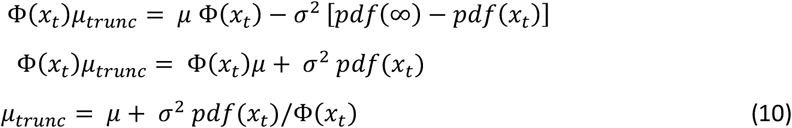

Later we will use the fractional truncation-selected change in mean, in units of initial *μ*:

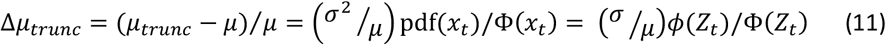

where *ϕϕ* is a sometimes-convenient form related to *pdf*(Methods).

*Δμ*_*trunc*_ assumes a Normal distribution, but utilizes no underlying system properties. Thus, Methods Eqn 11’s truncation-selected change, as for Methods Eqn 7 above, is general. Methods Eqn 11 is valid in any Normal system: genetic, pre-genetic, biological or chemical. Progress, *Δμ*_*trunc*_, will be proportional to coefficient of dispersion (simplified to “dispersion” in the main text), *σ/μ*, and to *ϕ*/Φ. The hatched blue region rightward in Fig. 1 illustrates a truncation selection.

### Truncation is somewhat enhanced by concurrent fitness

Threshold selection can also be somewhat more effective. Suppose that a characteristic being selected undergoes a truncation. For example, toleration of elevated temperature can aid survival, but a sudden temperature spike still truncates those beneath a new maximum. In Fig. 1, this selects the rightward blue hatched area, but now including individuals below the red dotted line. Below, such truncation on a selected background is quantitated.

We have already written the pdf for a fitness selection in Methods Eqn 4, and normalized it so that its range includes all valid *x* (Methods Eqn 6).

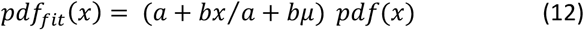

If we instead truncate the fitness distribution:

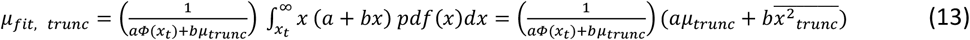

where 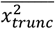 is the mean square *x* above the truncation. And 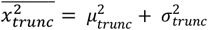 where, importantly, all quantities are for the initial Normal, and all are known. *μ*_*trunc*_ is Methods Eqn 10, and

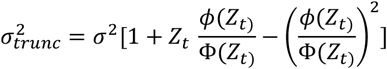

comes from standard sources (47). Combining all above as in Me*t*hods Eqn 13:

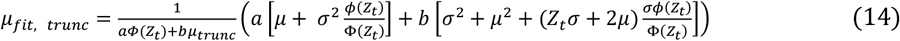

All quantities refer to the initial Normal, so are available via standard software. Moreover, combined selection and truncation in Methods Eqn 14, like separated fitness (Methods Eqn 6) and truncation (Methods Eqn 10) above, assume only properties of the distribution itself. The result is therefore again valid for any initially Normal system: protobiological, chemical or biological, independent of genetic or chemical detail.

### Numerical illustrations

Calculations were executed on a Dell XPS computer with an Intel Core i9-8950HK CPU @ 2.90GHz and 32 GB of RAM, running 64-bit Windows 10 Enterprise v. 20H2. Figures were prepared with Microsoft Excel 2016 16.0.5161.1002, making particular use of the Excel norm.dist function for Normal pdf, *ϕ*, and cumulative distributions, Φ.

As an example, in Methods Eqn 9, Φ*(x*_*t*_) is an integral not expressible in closed form using simple functions. But Φ*(x*_*t*_) is readily accessible via the widely-available cumulative Normal distribution function *C(μ, σ, x*_*t*_) (which yields the fraction of the distribution between -∞ and x_t_): Φ*(x*_*t*_*) = 1 - C(μ, σ, x*_*t*_).

### The Normal probability distribution function

The Normal probability distribution function used here appears in multiple forms:

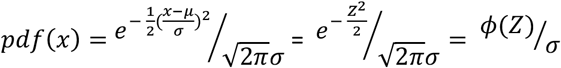

using text notation. The denominator’s standard deviation *σ* at right is significant because it differs from what is sometimes called the Standard Normal distribution *ϕ*, often seen in mathematical references (47). The Standard Normal has *μ* = 0, *σ* = 1, thus is not suited to investigation of variations. For example, the above denominator *σ* cancels an external factor of *σ*, clarifying *Δμ*_*trunc*_ dependence as ∝ *σ*, not *σ*^2^ (Methods Eqn 10, Fig. 3A). In the text, *pdf* and *ϕ* are interspersed where each was thought clearest.

## Data availability

All data analyzed during this study or required for its understanding are included in the manuscript.

